# Cognitive deficits found in a pro-inflammatory state are independent of ERK 1/2 signaling in the murine brain hippocampus treated with Shiga toxin 2 from enterohemorrhagic *Escherichia Coli*

**DOI:** 10.1101/2020.08.24.264929

**Authors:** Clara Berdasco, Alipio Pinto, Mariano Blake, Fernando Correa, Nadia A. Longo Carbajosa, Patricia Geogeghan, Adriana Cangelosi, Mariela M. Gironacci, Jorge Goldstein

## Abstract

Shiga toxin 2 (Stx2) from enterohemorrhagic *Escherichia coli* (EHEC) produces hemorrhagic colitis, hemolytic uremic syndrome (HUS) and acute encephalopathy. The mortality rate in HUS increases significantly when the central nervous system (CNS) is involved. Besides, EHEC also releases lipopolysaccharide (LPS). Many reports have described cognitive dysfunctions in HUS patients, the hippocampus being one of the brain areas targeted by EHEC infection. In this context, a translational murine model of encephalopathy was employed to establish the deleterious effects of Stx2 and the contribution of LPS in the hippocampus. Results demonstrate that systemic administration of a sublethal dose of Stx2 reduced memory index and produced depression like behavior, pro-inflammatory cytokine release and NF-kB activation independent of the ERK 1/2 signaling pathway. On the other hand, LPS activated NF-kB dependent on ERK 1/2 signaling pathway. Cotreatment of Stx2 with LPS aggravated the pathologic state, while dexamethasone treatment succeeded in preventing behavioral alterations. Our present work suggests that the use of drugs such as corticosteroids or NF-kB signaling inhibitors may serve as neuroprotectors from EHEC infection.

## Introduction

Shiga toxin 2 (Stx2) from Shiga toxin producing *Escherichia coli* (STEC) is one of the most virulent factors responsible for hemolytic uremic syndrome (HUS), a triad of clinical events that includes acute renal failure, nonimmune microangiopathic hemolytic anemia and thrombocytopenia. Prodromal bloody diarrhea may occur due to infection with diarrheagenic *Escherichia coli* pathotypes, mainly the serotype O157: H7 of enterohemorrhagic *Escherichia coli* (EHEC) [1–3].

The highest rates of HUS on the planet (more than 400 annual cases) are found in Argentina, where the incidence of pediatric cases reaches up to 17 cases per 100,000 children under 5 years of age [4]. The lethality reported in these patients is between 1 to 4% but it may rise up to 3-fold when the central nervous system (CNS) is involved [5–7].

Acute encephalopathy produced by STEC infection represents a critical episode in severe cases of disease [8]. We and other laboratories have demonstrated that Stx2 may exert a direct action in the CNS, in neurons and glial cells [9–12]. Furthermore, it has been reported that STEC may affect the CNS without renal involvement [13].

Stx2 produces its deleterious action through its canonical cell membrane receptor globotriaosylceramide (Gb3), a glycosphingolipid localized on the external bilayer lipid cell membrane. Once Stx2 binds to Gb3, the toxin reaches the ribosomes through a retrograde pathway and triggers ribotoxic stress leading to pro-apoptotic and pro-inflammatory mechanisms [14]. In addition to Stx2, the Gram-negative STEC also releases lipopolysaccharide (LPS), an endotoxin derived from its outer membrane [15]. It has been reported that patients with HUS/encephalopathy also present endotoxemia. Several animal models developed to study the effect of endotoxemia on STEC pathogenesis have shown that LPS amplifies the deleterious effects of STEC. In connection with this amplification, we have previously reported a relevant pro-inflammatory component in CNS damage produced by Stx2 [10, 11, 16].

Several reports have described cognitive dysfunction in HUS patients, the hippocampus being one of these damaged areas in the brain [17–20]. Alterations found in these patients include trouble finding words, severe alteration of consciousness, late memory decline, orientation deficits, seizures and coma. We have previously demonstrated morphological alterations induced by Stx2 and/or Stx2+LPS in the murine hippocampus [11]. In this regard, the aim of this work was to dig deeper in the understanding of the alterations mentioned above, which produce a cognitive decrease in memory and depression-like behavior related to hippocampal damage. Furthermore, we sought to establish whether pro-inflammatory involvement played a relevant role in pathogenesis and elucidate the intracellular mechanism likely to be involved in these events.

## Results

### Stx2+LPS reduced short term memory

The nose-poke habituation task was employed to establish whether the toxins change the capability of mice to store and evoke short term memories [24]. While no significant differences were found in exploratory activity among groups during TR (Control, LPS, Stx2 and Stx2+LPS), mice treated with Control, LPS or Stx2 yielded a significant reduction in the number of nose-pokes recorded between TR and TS (Fig 1B). These findings suggest that mice in these groups were able to store and retrieve short-term memory of the habituation task. Nevertheless, no significant differences were found between TR and TS for Stx2+LPS-treated mice, which is indicative of amnesia. Consequently, the memory index was significantly lower in Stx2+LPS-treated mice compared to all other treatments (Fig 1C) (p<0.05). No differences were observed in the memory index across Control, LPS and Stx2-treated mice.

**Figure 1.**
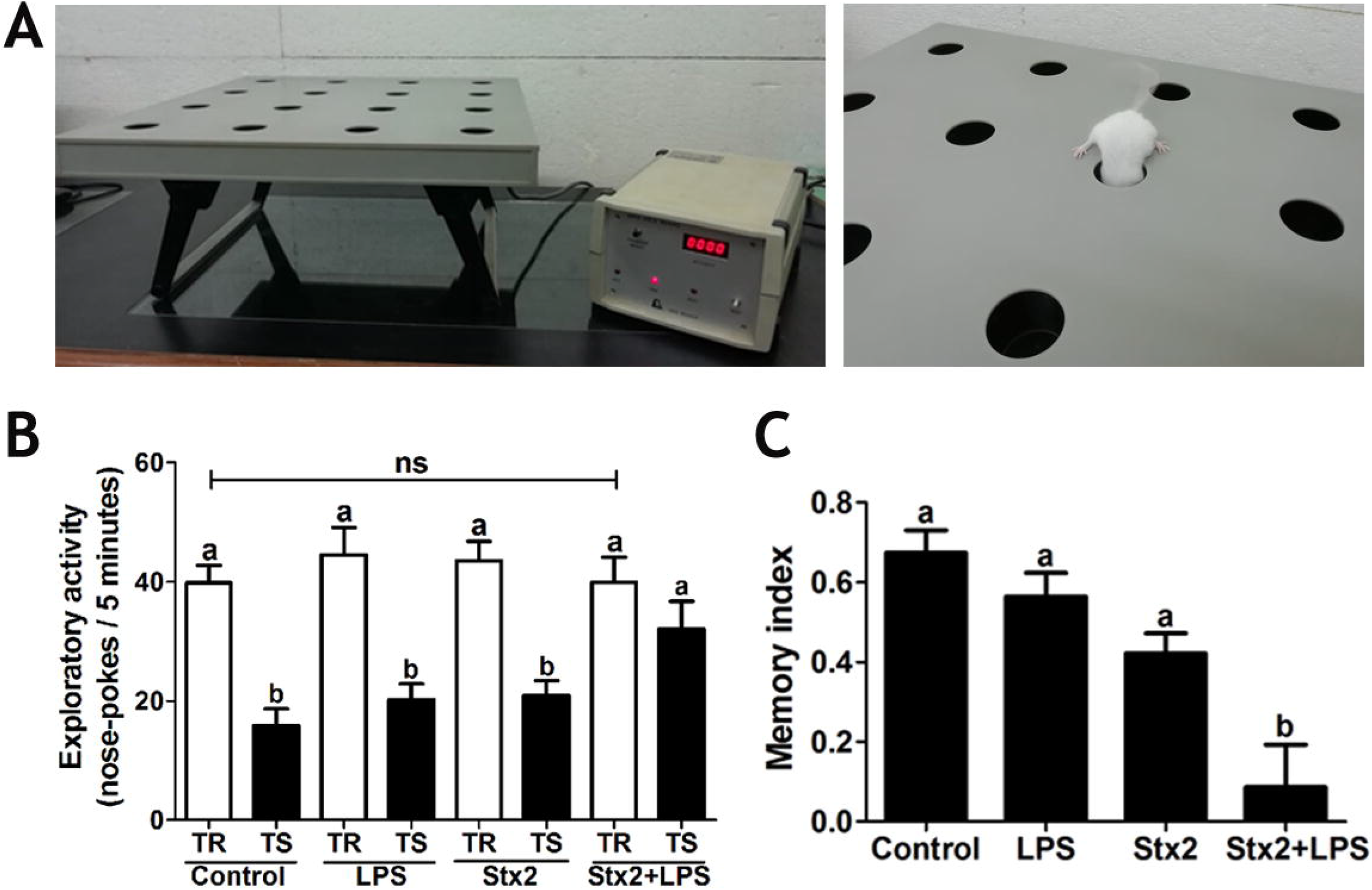
Short term memory. A: the hole-board apparatus (left), representative nose-poke assay (right). B and C: tests measured after 2 days of treatment. TR: training test; TS: retention test. Different letters (a, b) represent significant differences. Data were analyzed by non-parametric one-way ANOVA, Bonferroni’s *post hoc* test, p<0.05, n=15.

### Toxins produced MG reactivity in the CA1 hippocampal area

Iba1 is a microglial/macrophagic-specific calcium-binding protein which participates in membrane ruffling and phagocytosis in activated microglia [25]. An anti-Iba1 antibody was employed to establish whether the toxins produced activated MG morphology, as this molecule is upregulated in activated MG states [21, 22, 26].

After 2 days of treatment, the four MG morphological types were observed in the CA1 hippocampal area (Fig 2A). The MG activation score was significantly increased by both toxins compared to Control (Fig 2B), and maximally increased in Stx2+LPS-treated mice. No significant differences were found between LPS and Stx2-treated mice. In addition, the toxins also produced a significant increase in hyper-ramified, bushy and amoeboid MG activated morphologies as compared to Control (Fig 2C). In contrast, the toxins produced a significant decrease in ramified morphology (Fig 2C).

**Figure 2.**
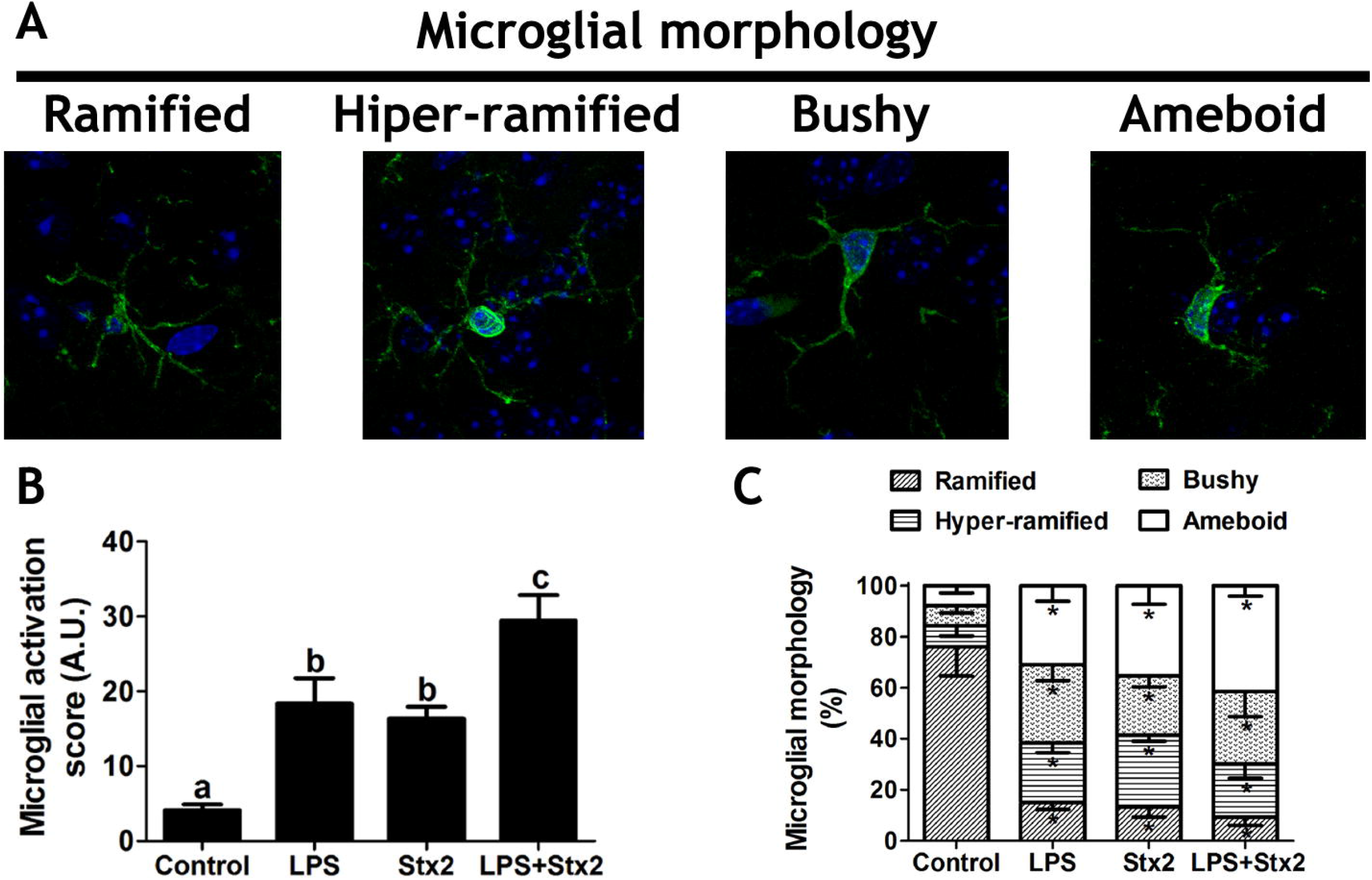
CA1 hippocampal MG response to toxins. A: Iba1 immunofluorescence of representative images from all distinct morphological MG types. B: microglial activation score after 2 days of treatment. Each letter (a, b, c) represents significant differences across each treatment. C: percentage of the microglial morphology in each treatment. Asterisks (*) represent significant differences across morphologies from all toxin-treated mice as compared to Control. Data from figures B and C were analyzed by one-way ANOVA, Bonferroni’s *post hoc* test, p<0.05, n=4.

### Toxins increased pro-inflammatory cytokines and decreased anti-inflammatory cytokines in the hippocampus

In order to establish whether these toxins altered the pro-inflammatory and anti-inflammatory cytokine profile in the mouse hippocampus treated with LPS and/or Stx2, an ELISA was performed to measure pro-inflammatory cytokines IL-6 and TNFα and anti-inflammatory cytokine IL-10. The expressions of IL-6 and TNFα were both significantly increased after 1 day of treatment in all toxin-treated mice (Fig 3A, B). On the other hand, IL-10 was significantly decreased after 1 day of treatment only in Stx2+LPS-treated mice (Fig 3C), but significantly decreased after 2 days of treatment in all toxin-treated mice (Fig 3D).

**Figure 3.**
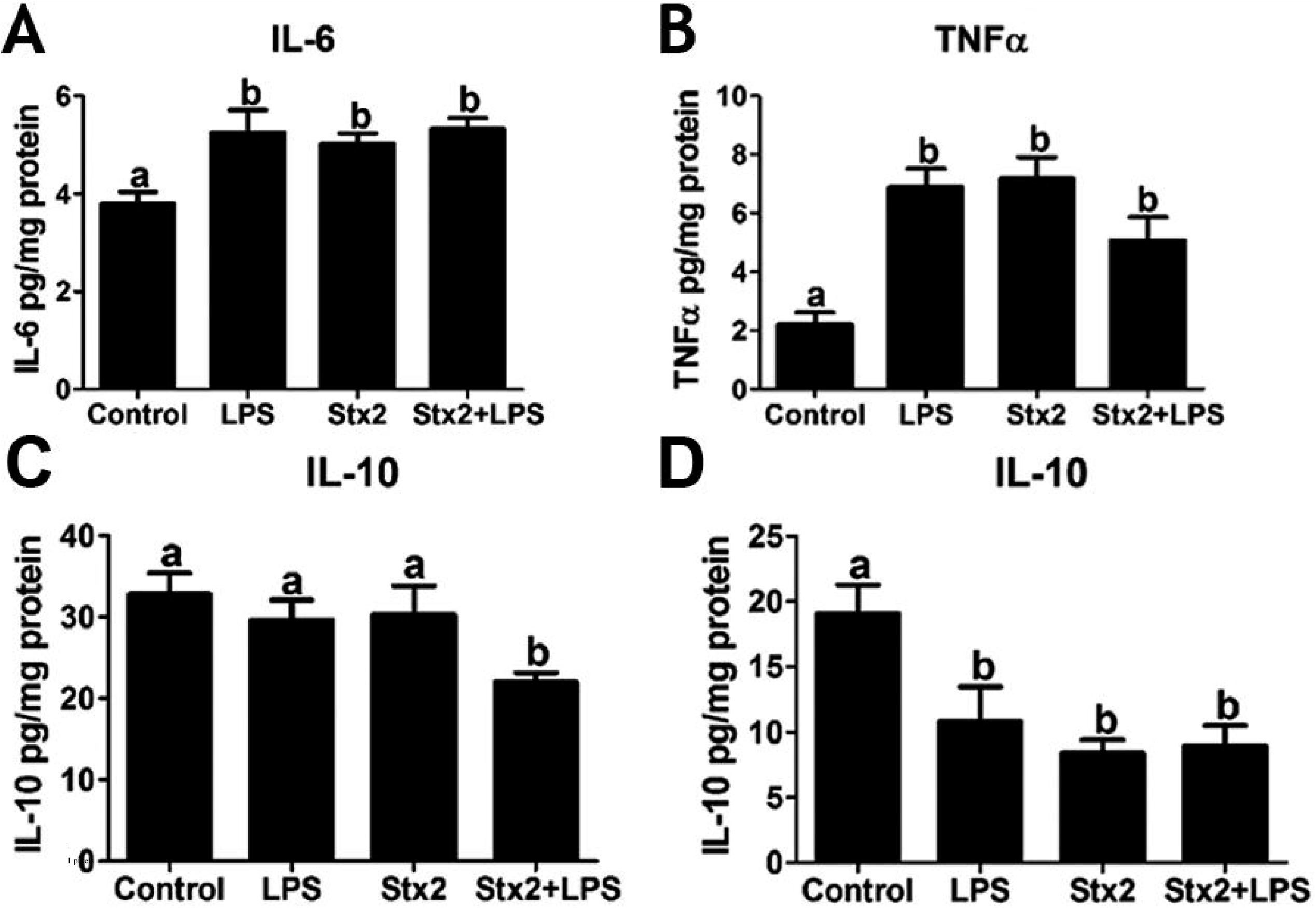
Pro- and anti-inflammatory cytokine measurements in the murine hippocampus. A: IL-6 after 1 day of treatment. B: TNFα after 1 day of treatment. C: IL-10 after 1 day of treatment. D: IL-10 after 2 days of treatment. Each letter (a, b) represents significant differences across treatments in all figures. Data were analyzed by one-way ANOVA, Bonferroni’s *post hoc* test, p<0.05, n=8.

### The pro-inflammatory profile induced by Stx2 was pIkB-dependent but ERK-independent

Given that toxins induced the expression of pro-inflammatory cytokines, we next studied the intracellular signaling involved in these processes. For this purpose, the ERK and IkB phosphorylation profiles were determined by Western blot following treatment with Stx2+LPS between 2 and 24 hours. As shown in figure 4A, upper panel, a significant reduction in phosphorylated ERK (pERK) was observed after 12 hours of treatment, while pIkB was increased after 2 hours (Fig 4B, upper panel).

**Figure 4.**
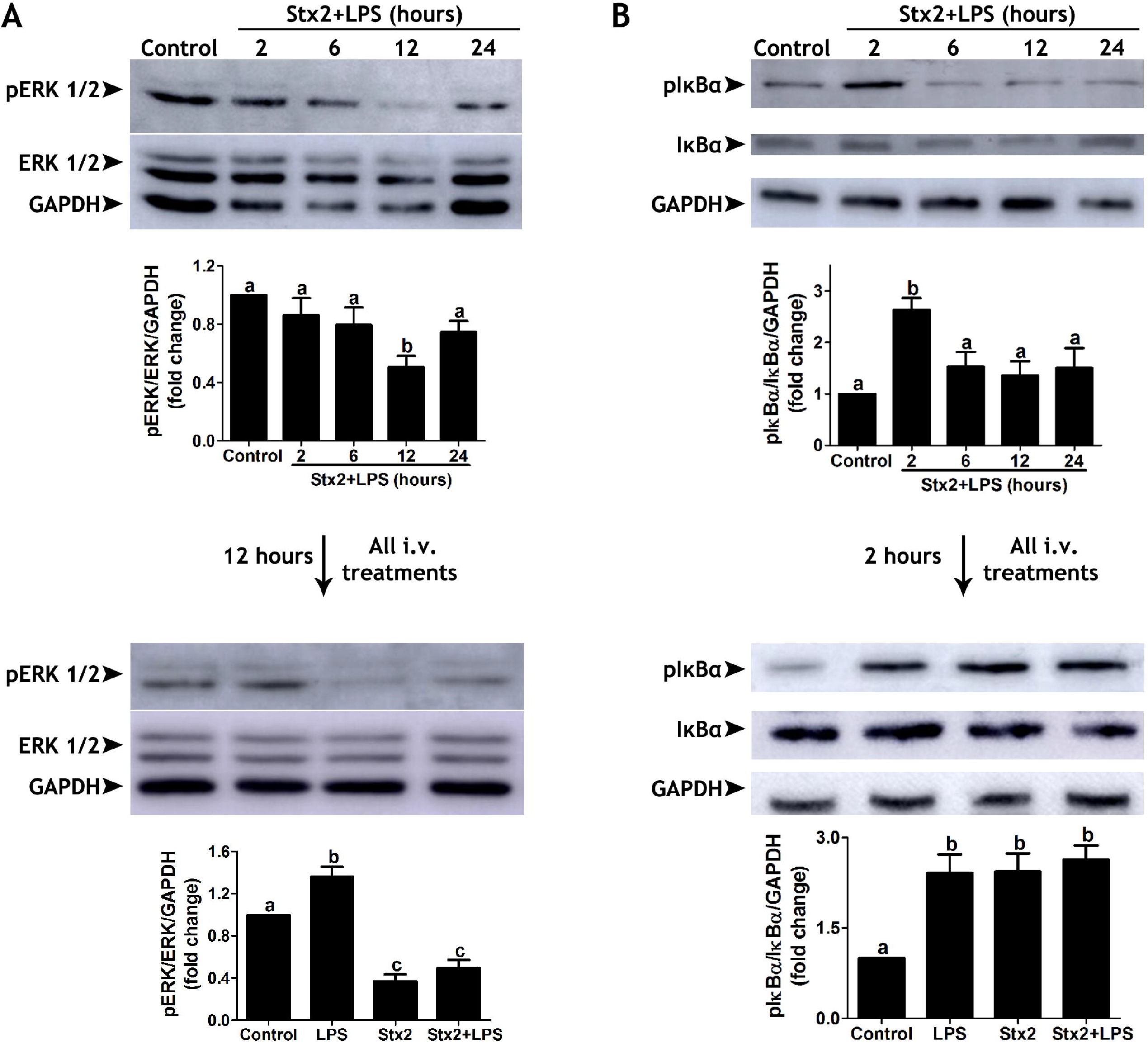
Changes in intracellular signaling. A, upper panel: Western blot of pERK 1/2 expression after 2, 6, 12 and 24 hours of Control or Stx2+LPS treatments; A, lower panel: Western blot of pERK 1/2 after 12 hours of the four treatments. B, upper panel: Western blot of pIĸBα after 2, 6, 12 and 24 hours of Control or Stx2+LPS treatments; B, lower panel: Western blot of pIĸBα after 2 hours of the four treatments. Each letter (a, b and c) represents significant differences across treatments. All data were analyzed by one-way ANOVA, Bonferroni’s *post hoc* test, p<0.05, n=8.

To investigate whether the response elicited by Stx2+LPS may be due to a synergistic effect of both compounds, we evaluated the individual effect of LPS or Stx2 on ERK and IkB phosphorylation. After 12 hours, Stx2 but not LPS alone induced a decrease in ERK phosphorylation (Fig 4A, lower panel), which suggests that the reduction in ERK activation induced by Stx2+LPS resulted from a predominant effect of Stx2. Regarding Ikb, 2 hours of treatment with Stx2 or LPS alone induced the same increase in IkB phosphorylation as combined Stx2+LPS treatment (Fig 4B, lower panel).

### Stx2 and Stx2+LPS induced depression-like behavior, which was attenuated by dexamethasone

To evaluate whether the toxins produced depression-like behavior and whether this behavior may involve an inflammatory component, toxin-treated mice were treated with corticosteroid dexamethasone and then subjected to the forced swimming test. Immobility time (considered when the animals were restricted to floating movements [23] was significantly higher in Stx2+LPS-treated mice as compared to Stx2, and also significantly increased in these two groups compared to LPS and Control (Fig 5). Treatment with dexamethasone significantly reduced immobility time in Stx2 and Stx2+LPS-treated mice (Fig 5). No significant differences were found between the i.p. treatments in Control and LPS-treated mice.

**Figure 5.**
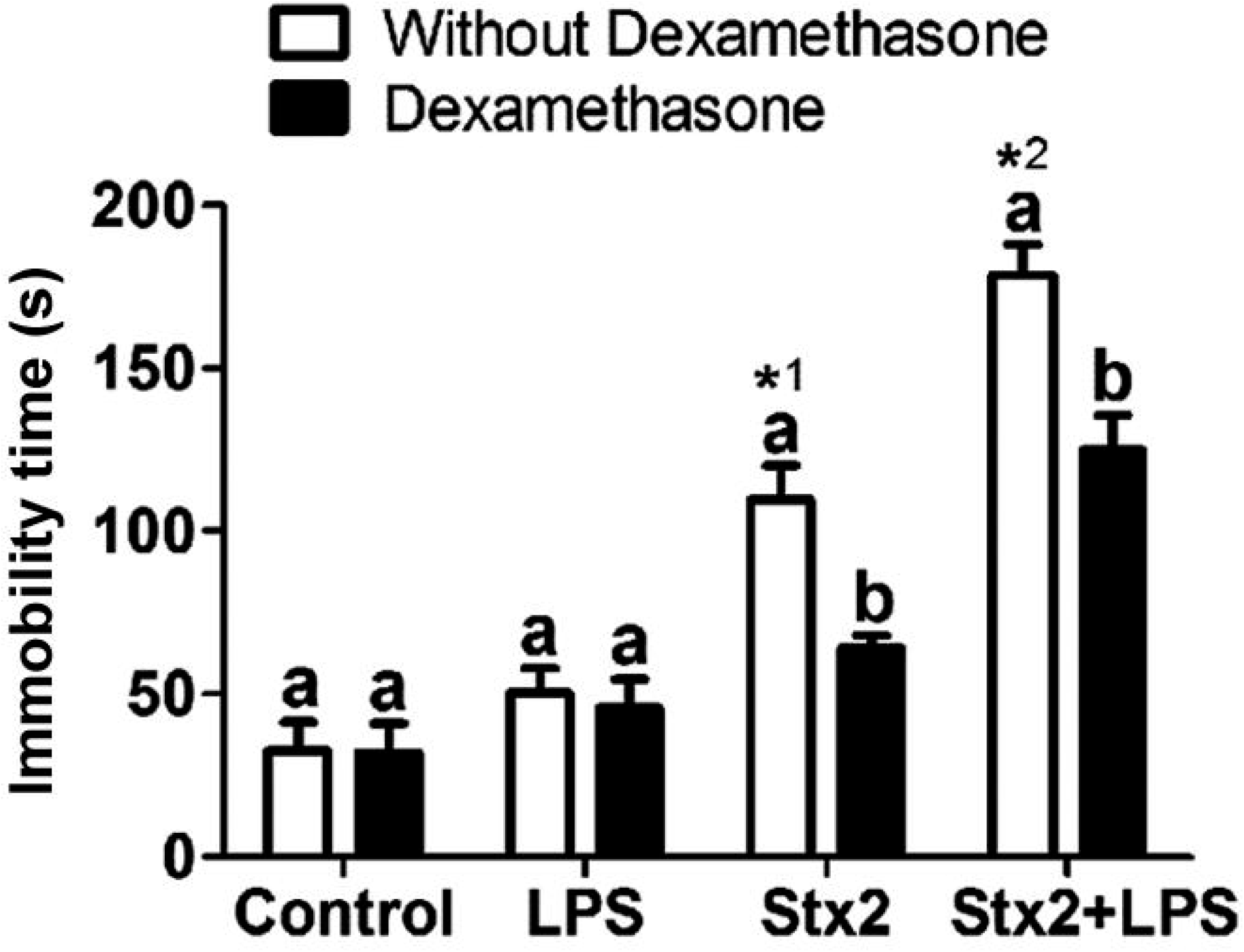
Depression-like behavior and the effects of dexamethasone. The effect of toxins and dexamethasone on mouse behavior after 2 days of treatment. Each letter (a, b) represents significant differences between i.p. treatments with or without dexamethasone. ^*1^ represents a significant difference between Stx2-treated mice without dexamethasone and the other treated mice without dexamethasone. ^*2^ represents significant differences between Stx2+LPS-treated mice without dexamethasone and the other treated mice without dexamethasone. Data were analyzed by non-parametric two-way ANOVA, Bonferroni’s *post hoc* test, p<0.05, n=12.

## Discussion

*In vivo* studies and clinical reports have shown the importance of neuroinflammation in the pathogenesis of Stx2-induced encephalopathy [10, 16, 27, 28]. However, none of these studies have shown the molecular pathways through which the inflammation is produced. The current report presents for the first time an introduction to the molecular mechanisms triggered by Stx2 in the murine brain hippocampus. Furthermore, it connects the inflammatory process triggered by Stx2 at the molecular level with behavioral changes, which confirms our previous results showing damage in the neurovascular unit produced by Stx2 in the CA1 hippocampal area [11].

The alterations in memory index and depression-like behavior produced by the toxins in the nose-poke habituation task and the forced swimming test, respectively, may be related with damage in hippocampal neurons and correlate with the cognitive imbalance reported in patients [17]. Considering previous reports [9–11], Stx2 may damage hippocampal neurons in at least two ways: first, Stx2 binds to the canonical Gb3 receptor localized in CA1 hippocampal neurons; and, second, indirectly by an inflammatory process with MG playing a fundamental role [10]. Indeed, we have recently reported in an *in vitro* model that Stx2 is up-taken by MG through its receptor Gb3, which produces an increase in MG metabolism, phagocytic capacity and pro-inflammatory cytokine release [29].

MG are mononuclear phagocytes resident in CNS parenchyma. MG are important sentinels responsible for CNS homeostasis which play an important role in synaptic organization, control of neuronal excitability, phagocytic removal of debris and trophic support leading to brain protection and repair [30, 31]. As primary innate immune cells, MG actively move through the CNS microenvironment while they sense disturbances in homeostasis by extending and retracting their fine processes [31]. Once they find some homeostatic imbalance through their wide range of receptors, MG change their morphology from a highly ramified small cell soma (traditionally called “resting”) to a variety of other distinct morphologies (traditionally called “activated”), ultimately becoming a poorly ramified amoeboid cell soma. These cells are also highly specialized in cytokine secretion [21, 22].

Stx2 and LPS produced a significant increase in the secretory MG morphology, with more amoeboid than ramified cells. Furthermore, this event was accompanied by a significant increase in pro-inflammatory IL-6 and TNFα and a significant reduction in anti-inflammatory IL-10 secretory profile. Traditionally, cytokine secretion has been classified according to its pro or anti-inflammatory profile. The M1 profile is responsible for the secretion of pro-inflammatory cytokines like IL-6 and TNFα. In contrast, the M2 profile is responsible for the secretion of anti-inflammatory cytokines such as IL-10 [32–34]. In addition, secretory ameboid MG morphology may also be an important oxidative source of reactive oxygen species (ROS) and nitric oxide (NO) radical derived products [35], as we have previously shown in the current model of Stx2-produced encephalopathy [11].

Cytokine synthesis is regulated by many intracellular signaling pathways triggered by different kinds of receptors that respond to different stimuli [36, 37]. For example, the binding of LPS to toll-like receptor 4 (TLR4) leads to the activation of the intracellular mitogen-activated protein kinase (MAPK) pathway and NF-κB nuclear translocation, which results in the secretion of pro-inflammatory cytokines [38, 39]. However, information regarding the intracellular signaling pathway triggered by Stx2 is very scarce, and particularly much less is known about it in brain parenchymal cells.

To our knowledge, all that is known so far about damage produced by Stx via the MAPK pathway is the activation of p38, which induces IL-8 secretion in the human colonic epithelial cell line [40], IL-1 secretion by human macrophages [41] and TNF production by a human adenocarcinoma-derived renal tubular epithelial cell line [42]. In addition, p38 activation has been also found partially responsible for Stx-induced cell death in Vero cells [43] and in intestinal epithelial cells [44]. On the other hand, the inhibition of p38 is a determinant of TNF up-regulation from Stx toxicity in brain microvascular endothelial cells [45]. Furthermore, signaling cascades including JNK and p38 molecules have been related to cell death by ribotoxic stress leading to apoptosis in intestinal epithelial cells [44].

MAPKs are a family of proteins which, in response to the activation of cell surface receptors, phosphorylate specific serine or threonine residues preceded by a proline residue on their substrates. Extracellular signals such as hormones, cytokines or growth factors activate the MAPK cascade which consists in a linear signaling triad comprising a MAPK kinase kinase (MAPKKK or MAP3K) and a MAPK kinase (MAPKK or MAP2K), ending in the phosphorylation and activation of MAPKs, which include the extracellular signal regulated kinases 1 and 2 (ERK1/2), p38 isoforms (α, β, γ, and δ) and c-Jun N-terminal kinases (JNK1, JNK2, and JNK3). The activation of MAPKs culminates with the phosphorylation of many cytosolic and nuclear substrates such as downstream kinases, phosphatases or transcription factors. The dysregulation of this intracellular cascade is implicated in many neurological disorders, including depression and Alzheimer’s disease [46]. Similar dysregulation was also found in our present work and could be compared to these neurological disorders.

According to the present data, in our encephalopathy murine model, the deleterious effect produced by Stx2 together with LPS [10, 11, 16, 29] suggests a potentially synergistic action rather than an additive one (Fig 6). While both toxins induced the activation and translocation of NF-κB to the nucleus, they performed it independently through two distinct pathways: LPS activated the ERK1/2 pathway, and Stx2 activated an ERK1/2-independent pathway. Furthermore, both pathways also produced the expression of genes related to a pro-inflammatory state such as TNFα and IL-6 and reduced the expression of those associated to anti-inflammatory cytokines such IL-10. Although further experiments are necessary to determine whether p38, JNK or a third unknown pathway may be regulated by Stx2 concerning NF-κB activation, it may be concluded that the activation of these molecular cascades may also be implicated in the activation of MG and concomitant cognitive deterioration observed in the current mouse model. Moreover, the fact that this deterioration has also been reported in patients suggests that the intracellular mechanisms studied in this work may also be at play in clinical scenarios. Since there is no consensus therapy currently available to treat STEC-derived encephalopathy, it is essential to elucidate the molecular mechanisms underlying the pathophysiological process of encephalopathy produced by Stx2 to propose therapeutic strategies which may efficiently protect the brain parenchyma [47]. Accordingly, our present work shows that the use of drugs such as dexamethasone or those blocking the cascade by preventing NF-kB translocation to the nucleus may serve as effective neuroprotectors with potentially beneficial use in the clinic.

**Figure 6:**
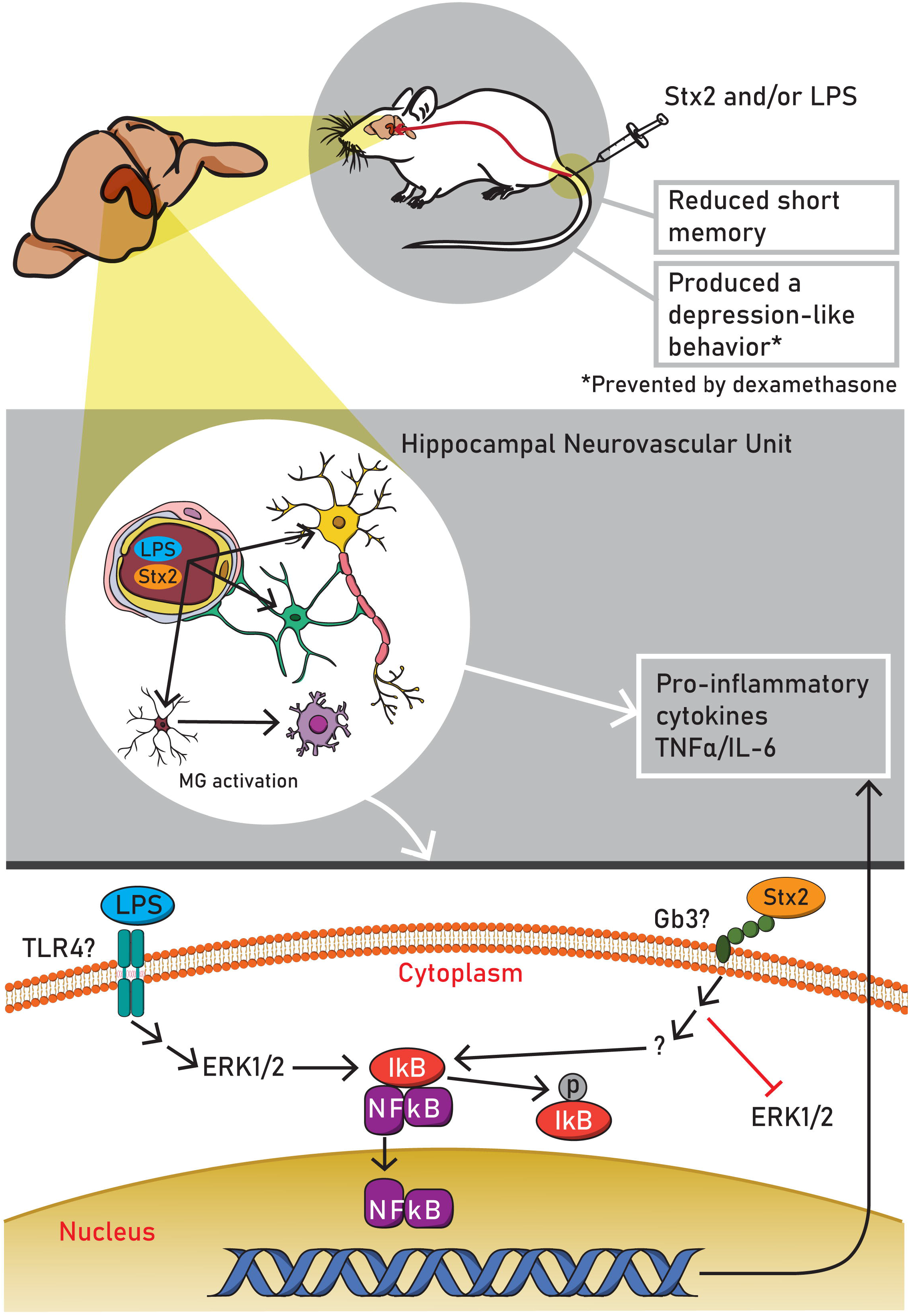
Suggested model of Stx2 and LPS intracellular deleterious actions in the murine hippocampus: The toxins reach the hippocampus crossing the blood-brain barrier that activate microglial cells. They activate NF-κB translocation to the nucleus through two distinct pathways engaging inflammatory processes and cognitive deficits. Astrocyte (green); MG cells (violet); neuron (yellow).

## Methods

### Animals

NIH male Swiss mice of 4 weeks (approximately 25±3 g) were housed under 12 h-light/12 h-dark cycles. Food and water were provided *ad libitum* and the experimental protocols and euthanasia procedures were reviewed and approved by the Institutional Animal Care and Use Committee of the School of Medicine, Universidad de Buenos Aires, Argentina (Resolution N° 046/2017). After an acclimatization period, animals were divided into four different groups according to intravenous (i.v.) treatment (the total amount of i.v. solution injected was 100 μl per mouse): Control (saline solution), LPS (800 ng), Stx2 (1 ng) and Stx2+LPS (1 ng and 800 ng, respectively). Stx2 dose was about 60% of the LD50 (1.6 ng per mouse). All the procedures were performed in accordance with the EEC guidelines for the care and use of experimental animals (EEC Council 86/609).

### Nose-poke habituation task

The apparatus employed was a hole-board (Ugo Basile Mod. 6650, Comerio, Italy) made of a matte gray Perspex panel (40 cm× 40 cm× 22 cm) which has 16 flush mounted tubes of 3 cm (Figure 1A). Each tube has an infrared emitter and a diametrically opposed receiver connected to an automatic counter to register the number of nose-pokes into the holes. The assay consisted of two stages: training (TR) and retention test (TS). TR was performed 2 days after the i.v. treatment. During TR, each mouse was placed at the center of the apparatus and the number of nose-pokes was automatically registered for 5 minutes (Fig 1A). During the TS stage, performed 30 minutes after TR, each mouse was placed again on the apparatus and the nose-pokes were registered for 5 minutes. The apparatus was carefully cleaned with ethanol 70% to avoid interference by previous animals’ performance. Exploratory activity was expressed as nose-pokes per 5 minutes. Differences in the number of nose-pokes between TR and TS represent the development of memory. A more significant difference between TR and TS means better memory formation. Results were also represented using a memory index derived from the following mathematical formula:

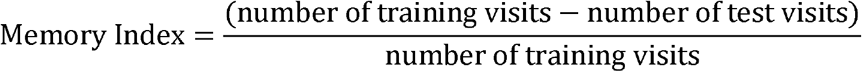

### Immunofluorescence assay

Two days after the i.v. treatments mice were anesthetized with sodium pentobarbital (100 mg/kg) and intracardiacally perfused with paraformaldehyde 4% diluted in 0.1 M phosphate buffer saline (PBS), pH 7.4. The brains were removed from the skulls, post-fixed overnight at 4 °C with the same fixative solution described above, cryopreserved daily with solutions with increasing concentrations of sucrose in PBS (10, 20 and 30%), and left overnight at 4 °C. Brain sections of 20 μm were obtained in a cryostat and stored at −20°C in a cryopreservation solution (50% PBS, 30% ethylene glycol and 20% glycerol) until the day of the immunofluorescence assay.

Immunofluorescence for ionized calcium binding adaptor molecule 1 (Iba1) was carried out to evaluate microglia (MG) reactivity. For this purpose, brain sections were blocked with a PBS solution containing 0.1% Triton X-100 and 10% goat serum to be incubated with a monoclonal goat anti-Iba1 antibody overnight at 4 °C (1:250 - Millipore, Temecula, CA, USA) followed by a donkey anti-goat Alexa Fluor 488 antibody (1:500 - Millipore, Temecula, CA, USA) for one hour at room temperature. All brain sections were also incubated with Hoechst 33342 (1:500 - Sigma, St. Louis, MO, USA) for 15 minutes at room temperature to detect brain cell nuclei. Negative controls were carried out by omitting the primary antibody. The hippocampal CA1 area (−1.70 and −1.82mm from bregma) was observed on an Olympus BX50 epifluorescence microscope equipped with a Cool-Snap digital camera and on an Olympus FV1000 confocal microscope.

Four distinct MG phenotypes immunolocalized in the hippocampal CA1 area were classified according to morphology [21]: ramified, hyper-ramified, bushy, and ameboid. A ramified morphology is associated with a resting microglial profile and was identified as a small soma with multiple long thin processes. The hyper-ramified phenotype was identified as a larger soma with thicker and branching processes. Bushy MG were also identified as having a larger soma with thicker but fewer processes. The ameboid morphology was identified as the largest soma with very few or absence of processes. The proportion of each type of morphology was measured for each treatment. Microglial activation was quantified using the following formula [22]:

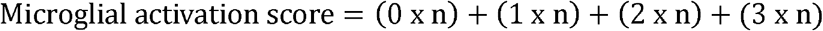

The numbers 0, 1, 2 and 3 represent arbitrary factors for the four different microglial morphologies (ramified=0, hyper-ramified=1, bushy=2, and amoeboid=3), and “n” represents the number of microglial cells with each specific morphology and corresponding arbitrary factor per micrograph.

### Enzyme-linked immunosorbent assay (ELISA)

Animals were sacrificed by decapitation 1 or 2 days after the four i.v. treatments, and the hippocampi were dissected and then homogenized on ice in PBS containing protease inhibitor cocktail (Sigma, St. Louis, MO, USA). Next, samples were centrifuged at 10,000 g for 15 minutes at 4 °C. IL-6, TNFα and IL-10 concentrations were immediately determined in the supernatant of murine hippocampal homogenates using the specific ELISA with antibodies and standards obtained from BD Biosciences (San Diego, CA, USA). The assay was performed according to the manufacturers’ instructions.

### Western blot

Animals were sacrificed by decapitation 2, 6, 12 and 24 hours after the i.v. treatments, and the hippocampi were dissected and homogenized in ice-cold lysis buffer (pH 7.4) containing 24 mmol/L HEPES, 1 mmol/L EDTA, 2 mmol/L tetrasodium pyrophosphate, 70 mmol/L sodium fluoride, 1 mmol/L β-glycerophosphate, 1% Triton X-100, 1 mmol/L PMSF, 10 μg/ml aprotinin and 2 μg/ml leupeptin (Sigma, St. Louis, MO, USA). Protein concentration in the homogenates was quantitatively determined through the Bradford assay. Hippocampal samples with equal amounts of protein (40 μg) were separated by electrophoresis in 10% sodium dodecyl sulfate polyacrylamide gels (SDS-PAGE) and transferred onto PVDF membrane (Bio-Rad Labs Inc.). Nonspecific binding sites on the membrane were blocked by incubation with 5% milk in Tris-buffered saline solution containing 0.1% Tween 20 at room temperature for 1 hour. The membranes were then incubated with primaries antibodies (1:500) overnight at 4 °C: rabbit anti-pERK1/2, rabbit anti-ERK1/2, rabbit anti-pIĸBα or rabbit anti-IĸBα. To avoid inaccuracies in protein loading, mouse anti-GAPDH was employed as an internal standard. All primary antibodies were purchased from Cell Signaling Technology, Inc. (Danvers, MA, USA). After 1-hour rinse, membranes were incubated with horseradish peroxidase-conjugated secondary antibody at 37 °C for 1 hour. Protein loading was evaluated by stripping and reblotting membranes with anti-GAPDH antibody (1:1000).

### Forced swim test

The four i.v. treated mouse groups described in *Animals* received either saline solution (without dexamethasone) or dexamethasone (7.5 mg/kg) by intraperitoneal (i.p. 100 μl) administration once a day for two consecutive days. Day zero was considered the day of the i.v. and i.p. treatments. Thus, the eight groups were divided as follows: Control with or without dexamethasone, LPS with or without dexamethasone, Stx2 with or without dexamethasone and Stx2+LPS with or without dexamethasone. After two days of treatments, each mouse was placed for 6 minutes in 3-liter beakers containing a volume of water at 25 °C that measured 15 cm height, which prevented mice from reaching the bottom of the container. The 6-minute test was recorded, and the time of immobility was measured only in the last 4 minutes [23]. After this time, each mouse was removed from the container and placed temporarily in a drying cage with a heat lamp above it. The water was changed after every session to avoid influence on the next mouse.

### Statistical analysis

All experimental data are presented as the mean ± SEM. Particularly, in the forced swim test, statistical significance was evaluated using two-way analysis of variance (ANOVA). For all other assays, statistical significance was evaluated using one-way ANOVA followed by Bonferroni’s *post hoc* test (GraphPad Prism 4, GraphPad Software Inc., San Diego, CA, USA). The criterion for significance was p<0.05 for all experiments.

## Acknowledgments

The authors are especially grateful to German Nicolás La Iacona for his technical assistance in using the confocal microscope. We are also grateful to María Marta Rancez for her special dedication and technical assistance.

